# AGZArank: Investigating epitope-conditioned antibody binder ranking with structure-derived synthetic supervision

**DOI:** 10.64898/2026.06.08.730711

**Authors:** Zhengisbek Sadykov, Aliya Khamidullina, Bolat Sultankulov, Dauken Seitkali

## Abstract

Computational antibody design methods can generate large libraries of candidate binders for a target epitope, but prioritizing which candidates to test experimentally remains a major bottleneck. Existing scoring approaches, including physics-based affinity estimators, structure-prediction-derived confidence measures, and inverse-folding likelihood models, provide useful proxy signals but are not explicitly optimized for early enrichment of binders among many structurally similar candidates.

Here we investigate epitope-conditioned antibody binder ranking as a dedicated learning problem and introduce AGZArank, a geometric deep learning framework trained with structure-derived synthetic supervision based on normalized pseudo-energy targets. On a benchmark of 45 experimentally validated antibody–antigen interfaces, AGZArank recovered the true binder within the top ten candidates in 44.4% of cases and showed stronger generalization on post-2021 structures than ProteinMPNN, ESM-IF, and PRODIGY. Ablation experiments indicate that ranking performance depends primarily on training scale and alignment between the optimization objective and retrieval-based evaluation, rather than architectural complexity alone. These results support candidate prioritization as a distinct and tractable problem in computational antibody design.

## 1 Introduction

Recent advances in generative protein design have made it possible to create large libraries of candidate antibody binders for a target epitope, using approaches such as inverse folding [1, 2], structure-conditioned sequence generation, and related geometric or language-model-based frameworks [3, 4]. As a result, the central bottleneck in many antibody design workflows is no longer candidate generation itself, but the ability to prioritize which candidates should be taken forward for experimental testing. In practical settings, only a small fraction of generated variants can be screened, so the critical requirement is not merely assigning a plausible compatibility score, but achieving strong early enrichment of the most promising binders within a ranked list.

A wide range of computational strategies have been explored for estimating antibody–antigen binding quality. Classical energy-based and statistical scoring approaches, including Rosetta-derived methods [5, 6], FoldX-like models [7], and related affinity predictors [8], provide physically interpretable estimates of interaction quality and have been widely used for mutation analysis and interface assessment. However, these methods typically depend on accurate docking poses, well-resolved side-chain geometries, and high-quality atomic structures. In generative design workflows, where candidate structures may be predicted, rebuilt, or only approximately placed relative to the target epitope, such assumptions are often violated, making direct affinity scoring difficult and unstable.

An alternative family of approaches uses protein structure prediction or complex prediction to infer binding compatibility indirectly. In principle, confidence measures from AlphaFold-like or docking-based pipelines [3, 9, 10] may correlate with successful interactions. In practice, however, these methods remain limited by the difficulty of accurately predicting antibody–antigen docking geometry, especially for flexible loops and subtle interface rearrangements. Small errors in docking position or side-chain placement can substantially affect downstream affinity estimates, reducing their usefulness for high-throughput candidate ranking.

Inverse folding models represent another important class of baselines. Methods such as ProteinMPNN [1] and ESM-IF [2] assign sequence likelihood conditioned on a fixed backbone structure and have become widely used in protein design because they effectively capture sequence–structure compatibility. Recent benchmarking studies [11, 12] have shown that such likelihood-based models can perform surprisingly well as enrichment baselines, and in some cases outperform classical physics-based approaches for identifying improved binders. At the same time, these models are not trained explicitly for binding ranking. Their scores reflect how plausible a sequence is for a given structure, rather than how well that sequence should be prioritized by binding potential relative to competing candidates. As a result, sequence likelihood is best understood as a strong but indirect proxy, not a ranking objective tailored to binder design.

More recently, there has been substantial progress in applying geometric deep learning and protein language models to antibody–antigen affinity prediction. EGNN-based models, graph neural networks, and hybrid architectures that integrate structural and sequence information [4, 13, 14, 15] have shown promising performance on mutation-level tasks such as ΔΔ*G* prediction. In parallel, synthetic-data frameworks have emerged as an effective way to overcome the limited scale and diversity of experimental affinity datasets. A recent example is Graphinity [16], which showed that large synthetic datasets combined with geometric deep learning can substantially improve antibody–antigen ΔΔ*G* prediction and support stronger generalization than experimental mutation data alone. These results highlight the importance of data scale and diversity for affinity modeling. However, Graphinity was trained and evaluated in a regression setting that lacks enrichment performance results.

Despite substantial progress in synthetic-data generation and affinity-prediction architectures, current approaches remain only partially aligned with the demands of practical binder design workflows, where the central requirement is not accurate regression of affinity changes, but early enrichment of the strongest binders among many plausible candidates. We argue that models optimized for regression accuracy do not necessarily produce the correct ordering of candidates, especially when the differences between variants are subtle and screening decisions depend on the top few ranked predictions. In other words, good affinity prediction and good candidate prioritization are related, but not equivalent. We therefore treat epitope-conditioned binder ranking as a distinct problem, in which success depends on robust prioritization under limited experimental validation capacity, imperfect structural inputs, and distribution shift.

Here, we investigate which factors most strongly determine ranking performance in this setting. In particular, we examine the role of the training supervision signal, the alignment between optimization objective and ranking metric, and the structural representation of the paratope–epitope interface. To study these questions, we train a paratope-focused EGNN [17] with ESM2 [18] embeddings as sequence features, using a structure-derived pseudo-energy target for supervision. We then compare ranking-aware and standard regression-based optimization to assess the contribution of objective alignment to ranking performance.

We evaluate these models using a controlled ranking benchmark built from experimentally observed antibody–antigen complexes paired with generated candidate variants, and assess performance by early-retrieval and enrichment metrics. We further test robustness on post-2021 structures, compare against representative likelihood-based and affinity-based baselines including ProteinMPNN [1], ESM-IF [2], and PRODIGY [8], and perform auxiliary mutation-level validation using AB-Bind [19]. Together, these experiments allow us to identify which components are necessary for effective and generalizable binder ranking in realistic antibody design workflows.

## 2 Results

We first provide an overview of the end-to-end candidate construction and ranking pipeline used throughout this study (Fig. 1). Candidate paratope sequences are converted into structural complexes via side-chain packing and subsequently evaluated by the AGZArank model, which predicts a normalized pseudo-energy used for ranking.

**Figure 1.**
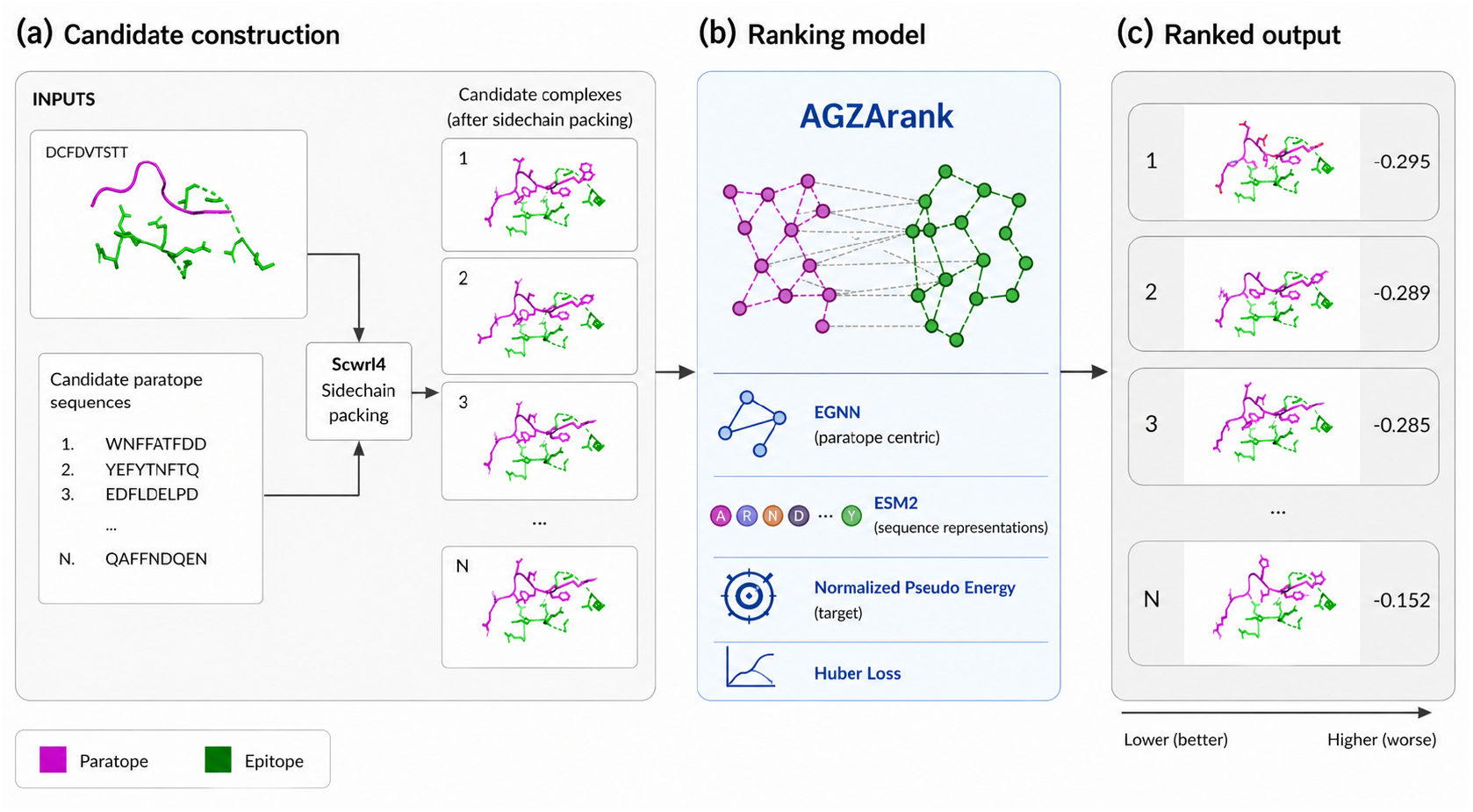
End-to-end antibody candidate ranking pipeline. **a**, Candidate construction. Input paratope sequences are converted to full-atom structures by packing onto fixed backbone templates with SCWRL4 [20], then combined with a fixed target epitope chain to generate candidate paratope–epitope complexes. **b**, Ranking model (AGZArank). Each complex is represented as a paratope–epitope interaction graph and processed using a paratope-centric equivariant graph neural network (EGNN) combined with sequence embeddings from ESM2. The model is trained to predict a normalized pseudo-energy using a weighted Huber loss. **c**, Ranked output. Candidate complexes are sorted according to predicted pseudo-energy, where lower values indicate more favorable interactions, enabling prioritization of top candidates for experimental validation.

### 2.1 Performance on the primary paratope ranking benchmark

We first evaluated whether the model can prioritize the experimentally observed binder among structurally plausible alternatives for the same target epitope. This setting reflects the intended use case in generative antibody design, where a large candidate library must be reduced to a small number of sequences for experimental validation. The benchmark comprised 45 independent paratope–epitope interface groups extracted from experimentally determined antibody–antigen complexes, each containing one experimentally observed paratope and 100 structurally consistent candidate variants scored using the learned pseudo-energy function.

The best-performing checkpoint was obtained at epoch 10. On the primary ranking benchmark, the model achieved a mean reciprocal rank (MRR) of 0.1764, an average rank of 32.7, and a median rank of 12. In parallel, regression performance against the structure-derived pseudo-energy target on the held-out validation set remained strong, with Pearson and Spearman correlations of 0.8438 and 0.8465, respectively.

Across the 45-group benchmark, the experimentally observed paratope was recovered within the top 10 ranked candidates in 44.4% of cases, within the top 5 in 31.1% of cases, and within the top 20 in 53.3% of cases. Thus, in nearly half of all benchmark groups, the true paratope was placed among the top 10 candidates out of 101 structurally similar alternatives.

These results indicate substantial early enrichment in the regime most relevant for experimental screening. Rather than merely producing reasonable continuous scores, the model places a large fraction of experimentally observed binders near the top of the ranked list, which is the central requirement for practical use in candidate prioritization workflows.

### 2.2 Comparison with inverse-folding baselines

We next asked whether the learned score provides an advantage over commonly used candidate-prioritization signals derived from inverse folding. For this comparison, we evaluated AGZArank against ProteinMPNN [1] and ESM-IF [2] on the same candidate pools, using sequence likelihood as the ranking signal for each baseline. We additionally included PRODIGY [8] as a representative physics-inspired affinity predictor, providing a complementary signal based on interface geometry and physicochemical properties rather than sequence likelihood. These models are strong comparators because they are widely used in protein design and often serve as practical enrichment baselines [11, 12].

On the full 45-group benchmark, AGZArank achieved stronger early-ranking performance than ESM-IF and was competitive with, or superior to, ProteinMPNN in the regime most relevant to screening. In particular, Top-10 recovery was 44.4% for AGZArank, compared with 40.0% for ProteinMPNN, 22.2% for ESM-IF, and 22.2% for PRODIGY.

Median rank was also markedly better than both ESM-IF and PRODIGY (12 versus 39 and 34, respectively), while ProteinMPNN showed somewhat stronger performance at larger *k* thresholds such as Top-20 and Top-50. A full summary of ranking performance on the complete candidate pools is provided in Table 1.

**Table 1.** Ranking performance on full candidate sets.

| Test set | Model | Top-$k$ recovery (%) | | | | | | Avg Rank | Median Rank |
| --- | --- | --- | --- | --- | --- | --- | --- | --- | --- |
|  |  | 1 | 3 | 5 | 10 | 20 | 50 |  |  |
| 45 PDB groups | AGZArank | 6.7 | 20.0 | 31.1 | 44.4 | 53.3 | 68.9 | 32.7 | 12 |
|  | ProteinMPNN | 8.9 | 24.4 | 26.7 | 40.0 | 60.0 | 77.8 | 27.0 | 17 |
|  | ESM-IF | 2.2 | 8.9 | 13.3 | 22.2 | 37.8 | 60.0 | 44.2 | 39 |
|  | PRODIGY | 6.7 | 11.1 | 13.3 | 22.2 | 37.8 | 57.8 | 43.0 | 34 |
| 15 PDB groups | AGZArank | 6.7 | 33.3 | 40.0 | 60.0 | 60.0 | 73.3 | 25.2 | 9 |
|  | ProteinMPNN | 0.0 | 6.7 | 6.7 | 13.3 | 46.7 | 80.0 | 33.0 | 23 |
|  | ESM-IF | 6.7 | 6.7 | 6.7 | 6.7 | 20.0 | 40.0 | 56.4 | 57 |
|  | PRODIGY | 13.3 | 20.0 | 26.7 | 33.3 | 60.0 | 73.3 | 27.3 | 20 |

The full candidate set includes the experimentally observed paratope and generated variants. Top- $k$ recovery denotes the fraction of interface groups in which the true binder is ranked within the top- $k$ candidates. The 15-PDB subset contains structures released after August 2021. Lower average and median ranks indicate better performance.

These results suggest that sequence likelihood is an informative but incomplete proxy for binder prioritization. ProteinMPNN retains useful enrichment, especially deeper in the ranked list, but its weaker precision in the earliest ranks indicates that structure compatibility alone does not reliably identify the strongest binders. This limitation is even more pronounced for ESM-IF, which showed consistently lower performance across early- and mid-ranking thresholds. PRODIGY, despite relying on physics-based interface features, showed moderate early enrichment but remained less effective than AGZArank in prioritizing the highest-quality binders.

Taken together, the full-benchmark comparison shows that the proposed scoring framework is not only effective in absolute terms, but also competitive with or superior to all baselines considered, with its clearest advantage appearing in early retrieval, where experimental decisions are typically made.

### 2.3 Performance under temporal distribution shift

A key concern for structure-conditioned baselines is whether apparent performance reflects dependence on historical structure–sequence regularities rather than transfer to genuinely newer interfaces. To address this, we repeated the evaluation on a temporally filtered subset consisting of 15 antibody–antigen complexes released after August 2021, while keeping candidate generation, scoring, and ranking metrics unchanged. This subset was designed as a stricter test of generalization to interfaces outside the training era of standard inverse-folding models.

On this post-2021 subset, AGZArank maintained strong early enrichment. Top-10 recovery reached 60.0%, with a median rank of 9. By contrast, ProteinMPNN achieved 13.3% Top-10 recovery with a median rank of 23, and ESM-IF achieved 6.7% Top-10 recovery with a median rank of 57 (Table 1). PRODIGY achieved 33.3% Top-10 recovery with a median rank of 20 on this subset, showing improved performance relative to the full benchmark but still lagging behind AGZArank.

The relative advantage of AGZArank became even clearer in this stricter setting than in the full benchmark. Both inverse-folding baselines showed substantial drops in early-ranking performance, whereas the learned energy-based score remained comparatively stable. PRODIGY showed improved absolute performance on this subset but still lagged behind AGZArank in Top-10 recovery, and its dependence on high-quality resolved structures limits its applicability in generative design workflows where candidate structures are predicted or rebuilt. This pattern is consistent with the view that explicit supervision on interface compatibility transfers more robustly to previously unseen antibody–antigen interfaces than sequence likelihood or physics-based scoring alone.

Although temporal filtering cannot eliminate every possible source of similarity, the magnitude and consistency of the performance gap support the conclusion that the proposed ranking signal generalizes more effectively under distribution shift than the baselines considered here.

### 2.4 Ranking performance depends on training scale and objective alignment

We next examined what most strongly determines ranking quality. Specifically, we asked whether performance arises mainly from architectural design, from limited training on experimentally determined interface fragments, or from the combination of large-scale synthetic supervision and an objective explicitly aligned with early enrichment.

To test the role of training scale and data regime, we compared the pretrained model with two matched-size alternatives: the same architecture trained from scratch on 608 interface samples derived from a non-redundant SAbDab set [21], and the same architecture trained from scratch on a randomly sampled synthetic subset of 608 training pairs. We also evaluated a fine-tuned model obtained by adapting the pretrained model on the same 608-sample SAbDab-derived set. In all cases, supervision was provided by the same structure-derived pseudo-energy target rather than by direct experimental affinity measurements.

Fine-tuning on the small SAbDab-derived set did not materially improve performance over the pretrained model. The fine-tuned model achieved an average rank of 32.76, a median rank of 12, and Top-10 recovery of 44.4%, effectively matching the base model. By contrast, both from-scratch small-data settings performed poorly: training on the 608-sample SAbDab-derived set yielded a median rank of 48 and Top-10 recovery of 11.1%, while training on the size-matched synthetic subset yielded a median rank of 55.0 and Top-10 recovery of 11.1%. These results, summarised in Table 2, indicate that ranking quality depends strongly on the scale and diversity of the full training distribution, rather than on limited exposure to experimentally determined structures alone. The lack of improvement after fine-tuning further suggests that the synthetic training corpus already captures most of the structural diversity needed for this task, and that the main limitation of the smaller datasets is insufficient coverage rather than data source.

**Table 2.**
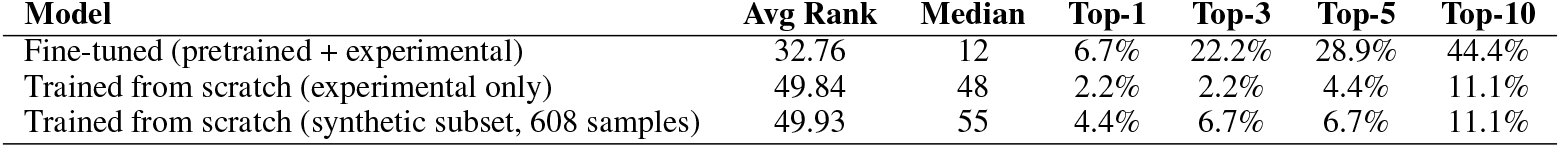
Comparison of ranking performance across training regimes with matched dataset size. Lower average and median rank indicate better performance, while higher Top-*k* values indicate improved retrieval of true binders.

| Model | Avg Rank | Median | Top-1 | Top-3 | Top-5 | Top-10 |
| --- | --- | --- | --- | --- | --- | --- |
| Fine-tuned (pretrained + experimental) | 32.76 | 12 | 6.7% | 22.2% | 28.9% | 44.4% |
| Trained from scratch (experimental only) | 49.84 | 48 | 2.2% | 2.2% | 4.4% | 11.1% |
| Trained from scratch (synthetic subset, 608 samples) | 49.93 | 55 | 4.4% | 6.7% | 6.7% | 11.1% |

We next tested objective alignment by replacing the percentile-aware weighted Huber loss with standard mean squared error. The weighted Huber objective was designed to give greater importance to lower-energy samples while remaining robust to outliers, thereby focusing learning on the part of the pseudo-energy distribution most relevant for early binder retrieval. Removing this emphasis produced the largest degradation observed in the study: Top-10 recovery fell from 44.4% to 17.8%, MRR dropped from 0.1764 to 0.0700, and Pearson and Spearman correlation with the structure-derived pseudo-energy target decreased to 0.5292 and 0.4882, respectively (Table 3).

**Table 3.** Ablation study on the 45-group ranking benchmark. Comparison of the full model with targeted modifications of individual components. All variants are trained on the same dataset with identical architecture, training protocol, and hyperparameters, unless otherwise specified. Lower average (Avg) and median (Med) rank indicate better performance; higher values are better for all other metrics.

| Model | Pearson | Spearman | MRR | Avg | Med | Top-1 | Top-3 | Top-5 | Top-10 |
| --- | --- | --- | --- | --- | --- | --- | --- | --- | --- |
| Full model | **0.8438** | 0.8465 | 0.1764 | 32.73 | **12.0** | 6.7% | **20.0%** | **31.1%** | **44.4%** |
| w/o ESM2 | 0.8425 | **0.8520** | **0.2070** | **29.51** | 20.0 | **13.3%** | **20.0%** | 22.2% | 33.3% |
| Vanilla EGNN | 0.8139 | 0.8004 | 0.1601 | 33.73 | 23.0 | 6.7% | **20.0%** | 22.2% | 33.3% |
| MSE loss | 0.5292 | 0.4882 | 0.0700 | 46.98 | 50.0 | 2.2% | 4.4% | 6.7% | 17.8% |

**Model definitions:** *Full model* — complete architecture including ESM2 embeddings, paratope-centric graph (no epitope–epitope edges), and percentile-aware weighted Huber loss.
*w/o ESM2* — sequence embeddings removed; node features include only amino acid, atom type, and node type embeddings.
*Vanilla EGNN* — graph includes epitope–epitope edges, restoring a standard fully connected interaction graph without paratope-centric restriction.
*MSE loss* — weighted Huber loss replaced with standard mean squared error (MSE) loss, with all other settings unchanged.

Together, these results show that effective paratope ranking depends primarily on broad structural coverage during training and on an optimization objective aligned with the strong-binding regime most relevant for downstream screening.

### 2.5 Architectural ablations identify contributions of sequence features and paratope-centric graph design

Having established that training scale and objective alignment are the dominant drivers of performance, we next asked which architectural choices provide additional benefit once these factors are in place. We therefore performed targeted ablations of the ESM2 sequence embeddings and the paratope-centric graph construction while keeping the remainder of the pipeline unchanged.

Removing ESM2 preserved strong regression behavior and even slightly increased Top-1 recovery and MRR, but reduced broader early-enrichment metrics. Specifically, the model without ESM2 achieved Top-5 and Top-10 recovery of 22.2% and 33.3%, compared with 31.1% and 44.4% for the full model, while median rank worsened from 12.0 to 20.0. This suggests that language-model-derived sequence information primarily improves ranking stability across a wider set of practically relevant top candidates rather than only the single highest-ranked position.

We also tested the contribution of the graph inductive bias by restoring epitope–epitope edges, thereby replacing the paratope-centric graph with a more generic interaction graph. This “vanilla EGNN” setting reduced performance, with MRR decreasing to 0.1601 and Top-10 recovery to 33.3%. These results support the view that restricting message passing to paratope–paratope and paratope–epitope interactions helps concentrate representation learning on interface-relevant signals. The full ablation results are reported in Table 3.

Overall, the ablation results indicate that the architecture does contribute meaningfully, but in a secondary role relative to training signal and objective alignment. Sequence-aware features improve broader early-ranking robustness, and paratope-focused graph construction improves interaction-specific representation learning.

### 2.6 External mutation-based validation on AB-Bind

To test whether the learned scoring function captures biologically meaningful affinity signals beyond the internal grouped benchmark, we evaluated it on a filtered subset of AB-Bind containing experimentally measured antibody–antigen affinity changes for point mutations. The analysis was restricted to entries with reported ΔΔ*G* values, heavy-chain mutations within CDR-H3, and complexes with at least five interacting epitope residues, yielding 96 variants in total, including 7 improved-affinity mutations defined by ΔΔ*G <* −0.5 kcal/mol. Variants were ranked by predicted pseudo-energy, and performance was assessed using enrichment metrics.

On this filtered subset, the model recovered 2 of the 7 improved-affinity variants within the top 5% of ranked candidates, corresponding to 28.6% recall and 5.49-fold enrichment. The same 2 improved variants remained within the top 10%, corresponding to 2.74-fold enrichment. These results, reported in Table 4, indicate that the learned score is sensitive to experimentally measured beneficial mutations and does not operate only within the synthetic or generative ranking setting.

**Table 4.** Enrichment of improved-affinity variants (ΔΔ*G < −*0.5 kcal/mol) on the filtered AB-Bind subset.

| Top-$k$ (%) | Tested | Tight found | Recall (%) | Enrichment |
| --- | --- | --- | --- | --- |
| 1 | 1 | 0 | 0.0 | 0.00 |
| 5 | 5 | 2 | 28.6 | 5.49 |
| 10 | 10 | 2 | 28.6 | 2.74 |
| 15 | 15 | 2 | 28.6 | 1.83 |

**Top- $k$ (%)**: fraction of ranked dataset considered. **Tested**: number of variants in this subset. **Tight found**: number of improved-affinity variants ( $\Delta\Delta G < -0.5$ ). **Recall**: percentage of all improved variants recovered. **Enrichment**: recall divided by dataset fraction.

As qualitative historical context, the original AB-Bind study reported enrichment values for classical scoring functions that were generally in the range of approximately 0.0–2.2 at the top 5% screening fraction and 0.4–2.9 at the top 10% fraction, depending on the method. Because our evaluation uses a substantially filtered subset tailored to the present model and input representation, these values should not be interpreted as a direct benchmark comparison. Nevertheless, the observed enrichment indicates that the learned ranking signal remains competitive in a challenging experimental point mutation-ranking setting.

Notably, recent synthetic-data affinity models such as Graphinity [16] have primarily been evaluated using correlation-based regression metrics rather than enrichment-oriented ranking measures. To examine whether strong regression performance necessarily translates into mutation-ranking utility, we derived AB-Bind-style enrichment values from Graphinity’s published prediction tables, as described in Methods. In that derived analysis on the full Graphinity test set (618 variants; 30 improved-affinity mutations under the same ΔΔ*G <* −0.5 threshold), enrichment was 0.665 at the top 5%, 1.661 at the top 10%, and 1.329 at the top 15%. Although this post hoc analysis is not directly comparable to the present filtered evaluation and was not reported in the original study, the resulting enrichment was modest relative to the regression performance emphasized in that work. This observation is consistent with our broader results, supporting the view that large-scale synthetic supervision alone is not sufficient for practical candidate prioritization unless the optimization objective is explicitly aligned with early retrieval.

Taken together, these results should be interpreted as supporting rather than definitive external evidence, given the limited number of improved-affinity variants after filtering. However, they strengthen the overall conclusion that the model captures interaction patterns with biological relevance beyond the internal benchmark, while also reinforcing the distinction between affinity regression and effective binder ranking.

### 2.7 Robustness to structural perturbation and candidate composition

Finally, we assessed robustness to structural perturbation and candidate composition in settings more reflective of fully generative antibody-design workflows. These analyses are important because practical design pipelines often rely on rebuilt, packed, or predicted candidate structures rather than experimentally resolved complexes, and because the experimentally observed binder is not available at inference time. For structural robustness, we applied controlled perturbations to the benchmark complexes, including local paratope rotation, translation along the epitope–paratope center-of-mass axis, atomic-coordinate jitter, and global complex rotation. Despite these perturbations, all ranking metrics remained unchanged, indicating that the learned score is stable to realistic geometric noise.

We also evaluated a stricter GenAI-only setting in which the experimentally observed paratope was removed from the candidate pool. In this regime, performance was measured by retrieval of the generated variant with highest sequence similarity to the experimental paratope. Although this setting is more challenging and represents a proxy rather than exact binder recovery, AGZArank retained meaningful enrichment, achieving 37.8% Top-10 and 75.6% Top-50 recovery on the full 45-group benchmark, and 53.3% Top-10 recovery on the post-2021 subset (Table 5). PRODIGY also showed improved performance relative to its full-candidate setting, but remained less consistent across ranking thresholds. Together, these results support the practical robustness of the ranking framework under imperfect structures and fully generated candidate sets.

**Table 5.** Ranking performance on generated candidate sets.

| Test set | Model | Top-*k* recovery (%) |  |  |  |  |  | Avg | Median |
| --- | --- | --- | --- | --- | --- | --- | --- | --- | --- |
|  |  | 1 | 3 | 5 | 10 | 20 | 50 | Rank | Rank |
| 45 PDB groups | AGZArank | 4.4 | 13.3 | 17.8 | 37.8 | 51.1 | 75.6 | 29.9 | 19 |
|  | ProteinMPNN | 11.1 | 15.6 | 24.4 | 42.2 | 55.6 | 86.7 | 23.0 | 14 |
|  | ESM-IF | 11.1 | 13.3 | 17.8 | 35.6 | 55.6 | 77.8 | 27.2 | 18 |
|  | PRODIGY | 2.2 | 13.3 | 13.3 | 24.4 | 33.3 | 64.4 | 37.7 | 28 |
| 15 PDB groups | AGZArank | 6.7 | 13.3 | 20.0 | 53.3 | 53.3 | 80.0 | 27.6 | 9 |
|  | ProteinMPNN | 13.3 | 13.3 | 26.7 | 40.0 | 53.3 | 86.7 | 24.0 | 20 |
|  | ESM-IF | 6.7 | 13.3 | 13.3 | 26.7 | 53.3 | 73.3 | 30.6 | 19 |
|  | PRODIGY | 6.7 | 20.0 | 20.0 | 26.7 | 33.3 | 66.7 | 41.3 | 28 |

In the GenAI-only setting, the experimentally observed paratope is removed from the candidate pool. Performance measures retrieval of the generated variant most similar to the experimental binder. Top-*k* recovery denotes the fraction of interface groups in which this target variant is ranked within the top-*k* candidates. Lower average and median ranks indicate better performance.

## Discussion

Prioritizing binders to a target epitope remains a central challenge in computational antibody design, particularly in screening settings where only a small number of generated candidates can be tested experimentally. Existing computational signals are informative but indirect: structure prediction and docking provide candidate poses and proxy confidence signals, inverse-folding models assess sequence–structure compatibility, and most machine-learning approaches focus on mutation-level affinity prediction. None of these formulations is designed specifically for early retrieval of the best binders in a de novo ranking setting.

AGZArank was developed to address this gap. The model combines a paratope-centric EGNN [17] with ESM2 [18] sequence embeddings, trained on a large-scale synthetic dataset using a structure-derived normalized pseudo-energy target and a weighted Huber loss designed for early binder enrichment. Because no standard benchmark exists for de novo paratope ranking at a fixed epitope interface, we also constructed a benchmark tailored to this use case, using experimentally observed paratope–epitope interface groups paired with epitope-conditioned generated paratope candidates. Although retrospective, this design reflects the practical decision regime of screening pipelines more directly than mutation-regression datasets.

Under this evaluation, AGZArank showed strong early enrichment on the developed benchmark and retained the strongest performance under temporal distribution shift. On structures released after August 2021, AGZArank achieved 60.0% Top-10 recovery, compared with 13.3% for ProteinMPNN [1], 6.7% for ESM-IF [2] and 33.3% for PRODIGY [8]. The widening gap under temporal filtering supports the view that the learned ranking signal transfers more robustly to previously unseen interfaces than inverse-folding likelihood or affinity-oriented structure-based scoring alone. More broadly, these results indicate that binder prioritization is not well captured by sequence compatibility or generic affinity-scoring proxies, but should instead be treated as a distinct learning problem.

The strongest mechanistic evidence for this conclusion comes from the loss-function ablation. Replacing the percentile-aware weighted Huber objective with mean squared error reduced Top-10 recovery from 44.4% to 17.8% and more than halved mean reciprocal rank, a larger degradation than that produced by any architectural modification. This shows that, in screening-oriented pipelines, alignment between the optimization objective and the retrieval task can be at least as important as model architecture. The ablation study also showed that restricting message passing away from epitope–epitope edges improved performance relative to a more standard graph construction, supporting the value of interface-focused modeling in this asymmetric setting. ESM2 [18] embeddings further improved ranking stability across top candidates. Taken together, these results suggest that architectural choices matter, but their effect is smaller than that of objective alignment.

The training-regime comparisons reinforce this interpretation. Training from scratch on small datasets performed poorly whether the structures came from a non-redundant SAbDab [21] set or from a size-matched synthetic subset, and fine-tuning on experimentally determined interfaces did not improve performance over the pretrained model. These findings suggest that robust generalization depends on broad structural coverage and that the synthetic corpus already captures much of the diversity required for the task. At the same time, these results suggest that corpus scale alone is unlikely to account for the observed gains. Because matched-size training from scratch remained weak whereas the full pretrained model generalized substantially better, the structure-derived normalized pseudo-energy target likely serves as a useful organizing signal for ranking. A plausible interpretation is that normalization improves comparability across candidates rather than allowing scores to scale primarily with interface size. Together, broad structural coverage and normalized structure-derived supervision appear to provide an effective training signal for ranking. This is practically important because such supervision can be assembled at scale without the throughput constraints of direct binding measurement.

The auxiliary validations further reinforce the distinction between regression and retrieval. On a filtered AB-Bind [19] subset, AGZArank enriched experimentally improved-affinity mutations above random expectation, indicating that the learned score captures biologically meaningful signal beyond the synthetic and generative settings in which it was trained. At the same time, enrichment values derived from Graphinity’s [16] published predictions suggest that strong mutation-level regression performance does not necessarily translate into strong early enrichment. These results support the view that regression and ranking are related objectives, but not interchangeable ones.

Several limitations define the scope of these conclusions. The framework assumes a structurally specified local binding interface and is therefore most applicable when a valid paratope–epitope arrangement is already known or fixed and the task is to rank designed or mutated loop sequences on that interface. It is not directly applicable when only antibody sequence is available or when no reliable docking configuration has been established. In addition, the primary benchmark is retrospective, and further prospective and experimental validation will be important for assessing how well the observed ranking gains translate to real screening campaigns.

Overall, this study supports a clear conclusion: effective candidate prioritization depends on aligning supervision and optimization with the retrieval criteria that govern experimental screening. More broadly, ranking in computational design should be treated as a problem in its own right rather than as a by-product of affinity regression.

## 4 Methodology

### 4.1 Task formulation

We formulate antibody–antigen binder ranking as an epitope-conditioned paratope scoring problem. Given the atomic coordinates of a target epitope together with a candidate paratope (specified by backbone geometry and amino-acid sequence, which is subsequently realized into a full-atom structure), the model outputs a scalar normalized pseudoenergy that quantifies interface compatibility.

Although the model is trained as a regression problem against this structure-derived target, performance is evaluated primarily in terms of group-wise ranking and early enrichment: for each of 45 experimentally validated antibody– antigen complexes, the model must recover the true binder among 100 structurally consistent decoy variants generated for the same epitope.

This formulation explicitly aligns model optimization with the practical requirements of generative antibody design pipelines, where only a small number of top-ranked candidates can be experimentally tested.

### 4.2 Training data

We constructed a proprietary synthetic dataset of approximately 45,000 paratope–epitope-like structural pairs derived from experimentally determined protein structures. Each sample consists of a short interacting fragment represented at full atomic detail using the ATOM14 scheme [22], where up to 14 heavy atoms are defined per residue and missing atoms are masked.

To ensure compatibility with generative design pipelines, which typically provide backbone geometry or aminoacid sequences without explicit side-chain coordinates, all structures underwent rotamer library-based side-chain reconstruction using SCWRL4 [20]. During training, geometric augmentations were applied to the input complexes while the normalized pseudo-energy target remained fixed to the original, unperturbed geometry. Specifically, the paratope was subjected to a local random rigid-body rotation about the epitope center of mass (up to 15^*°*^) and a translation along the epitope–paratope center-of-mass axis (up to 1.5 Å), followed by Gaussian jitter of real atomic coordinates (*σ* = 0.05 Å) and an additional random global rotation of the full complex (up to 180^*°*^). Placeholder atoms encoded as zero coordinates were excluded from perturbation. This augmentation strategy simulates docking uncertainty and coordinate noise commonly encountered in candidate generation, while preserving a consistent supervision target defined on the original interface geometry.

The dataset was constructed independently of the experimentally observed antibody–antigen complexes used in the evaluation benchmark, so that evaluation reflects generalization to previously unseen interfaces rather than overlap with training data.

### 4.3 Pseudo-energy target

For each paratope–epitope complex in the training set, a normalized pseudo-energy describing interface compatibility is computed directly from atomic geometry and used as the regression target. Rather than using raw interaction sums, we train against a contact-normalized pseudo-energy that provides a more consistent measure of interface compatibility across fragments of varying size.

#### 4.3.1 Interaction filtering

To restrict computations to physically relevant interactions, a two-stage filtering procedure is used. First, residue pairs are considered only if the distance between their C_*α*_ atoms satisfies

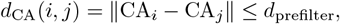

ensuring interface locality. Second, atom pairs are evaluated only if their interatomic distance

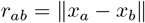

is smaller than a maximum interaction distance *r*_max_. This procedure restricts interactions to the local interface region while maintaining smooth dependence on atomic geometry.

To avoid discontinuities associated with hard contact cutoffs, pairwise interactions are weighted using a smooth contact kernel

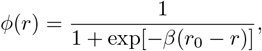

which provides continuous interaction weights and improves robustness to small structural perturbations. To prevent divergence of short-range interaction terms, interatomic distances are clamped

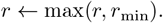

#### 4.3.2 Pairwise interaction terms

For each atom pair, three physics-inspired interaction components are evaluated.

##### Van der Waals interaction

which captures steric packing and dispersion interactions based on atom-type parameters.

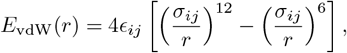

##### Electrostatic interaction

where *q*_*i*_ and *q*_*j*_ denote atom charges and *ε*(*r*) is a distance-dependent dielectric.

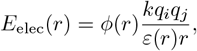

##### Non-polar interaction

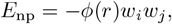

where *w*_*i*_ and *w*_*j*_ represent atom-level hydrophobicity weights.

The total interface energy is obtained by summing these contributions over all interacting atom pairs:

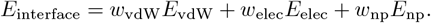

#### 4.3.3 Interface normalization

Direct summation of pairwise interaction terms produces interface scores that are strongly confounded by interface size, because larger paratope–epitope regions naturally accumulate more interaction contributions regardless of interaction quality. Normalization is therefore required to make pseudo-energy values comparable across candidate complexes with different effective interface extents. Rather than dividing by raw atom count or a hard contact count, we normalize by an effective contact mass derived from a continuous smooth contact kernel that weights atomic interactions by proximity without a binary distance cutoff. Specifically, because larger interfaces naturally accumulate more pairwise interactions, the raw pseudo-energy is normalized by the effective contact mass

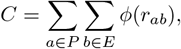

yielding the final normalized pseudo-energy

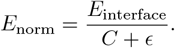

Negative values indicate favorable interfaces, while positive values correspond to unfavorable or non-complementary interactions.

### 4.4 Model Architecture

To model antibody–antigen interactions, we adopt a hybrid architecture that combines geometric reasoning with sequence-derived representations. The structural component is implemented using an equivariant graph neural network (EGNN), while evolutionary information is incorporated via embeddings from the ESM2 protein language model. An overview of the framework is shown in Fig. 3.

**Figure 2.**
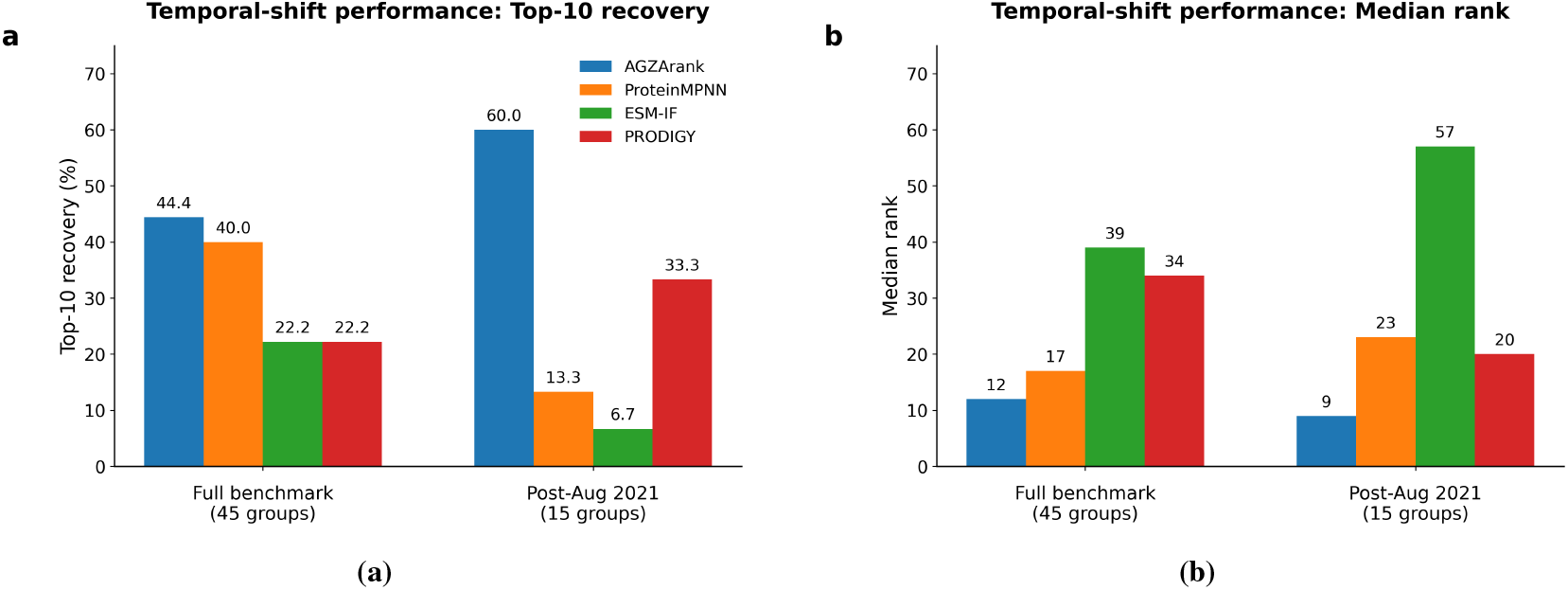
Ranking performance on the full benchmark and a temporally filtered subset. **a**, Top-10 recovery and **b**, median rank for AGZArank, ProteinMPNN, ESM-IF, and PRODIGY on the full 45-group benchmark and the post-August 2021 subset of 15 paratope–epitope interface groups. Temporal filtering was used to create a stricter generalization setting outside the likely training era of standard inverse-folding baselines.

**Figure 3.**
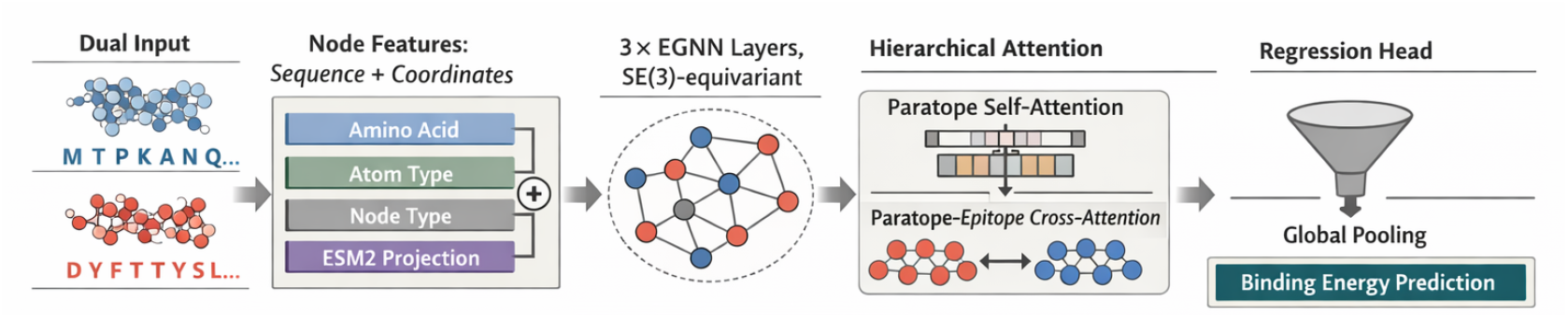
End-to-end paratope-focused geometric deep learning framework for antibody–antigen binding energy prediction. Epitope and paratope atomic coordinates are centered and represented as graph nodes with atom-type, residue-type, and node-type features. Sequence embeddings derived from ESM2 are projected into the structural latent space and fused with atomic representations prior to equivariant message passing. A paratope-centric EGNN updates node features while preserving coordinate equivariance. Hierarchical attention aggregates atom-level embeddings into residue representations, followed by paratope self-attention and cross-attention between paratope and epitope residues to model interface-specific interactions. Global pooling produces a compact complex-level representation used by the regression head to predict binding energy.

#### Input features

Atomic coordinates and graph-based features are processed at the atom level. Each node is described by four feature types: amino-acid identity, atom type, node type (epitope or paratope), and projected ESM2 [18] residue embeddings. All features are embedded into a shared 128-dimensional latent space and combined via elementwise summation, encouraging alignment between sequence-derived and structure-derived representations. Atomic coordinates are centered by subtracting the complex center of mass to ensure translational invariance.

#### Paratope-centric graph construction

Graphs are constructed using a 6 Å distance threshold for initial edge definition. Edges are categorized as paratope–paratope, epitope–epitope, or inter-chain (paratope–epitope). Interatomic distances are encoded using 32 Gaussian radial basis functions, and edge types are represented with one-hot encodings. During message passing, neighborhoods are further restricted to a maximum interaction radius of 8 Å and at most 16 nearest neighbors. Epitope–epitope edges are explicitly removed, restricting information flow to paratope–paratope and paratope–epitope interactions only. This paratope-centric design biases representation learning toward interface-relevant signals while avoiding dilution by intra-epitope correlations. The contribution of this inductive bias is evaluated in ablation studies (Section 2.5).

#### EGNN blocks

Node features are processed through three equivariant message-passing layers implemented with EGNN [17]. Each layer preserves SE(3) equivariance while updating only node features (update_coors=False), keeping atomic coordinates fixed to prevent structural drift. Message passing operates on the local neighborhood defined above, with interatomic distances encoded using Fourier features (eight frequencies). The resulting atom-level representations are projected into the downstream latent space.

#### Hierarchical attention

Instead of simple mean pooling, we introduce a learnable atom-to-residue attention mechanism. A linear scoring layer produces per-atom attention weights that are normalized via softmax, yielding residue representations as weighted sums of atomic embeddings.

Paratope residues are further refined using a two-layer transformer encoder [23] (four attention heads) to capture intra-paratope dependencies. Cross-attention is then applied with paratope embeddings as queries and epitope embeddings as keys/values, followed by reverse cross-attention to enable bidirectional interface conditioning. This hierarchical attention mechanism improves sensitivity to both structural and sequence variation.

#### Output head

Global pooling is performed separately on the updated paratope and epitope residue representations using learnable attention weights. The resulting complex-level vectors are concatenated and passed through a three-layer MLP regression head (with dropout) to predict the normalized pseudo-energy *E*_norm_. The model is trained end-to-end using the percentile-aware weighted Huber loss described below.

### 4.5 Optimization

The model is trained to predict the normalized pseudo-energy *E*_norm_ using a regression objective designed to emphasize the part of the target distribution most relevant for candidate prioritization. In the downstream ranking task, the key requirement is not equally accurate ordering across all complexes, but reliable identification of the lowest-energy candidates, which correspond to the strongest putative binders. Errors among high-energy candidates are therefore less consequential than errors that cause strong binders to be misplaced or lost from the top of the ranked list.

To reflect this asymmetry, we use a percentile-aware weighted Huber loss [24]. Given predicted energy *ŷ*∗ *i* and ground-truth energy *y*_*i*_, samples with lower energies are upweighted according to their position relative to the 40th percentile of the training distribution (*p ∗* 40). The weighting function is defined as:

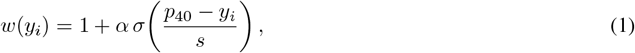

where *σ* (·) denotes the sigmoid function, *α* controls the strength of upweighting, and *s* determines the smoothness of the transition.

The final objective is:

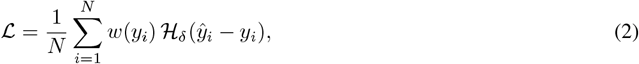

where *H*_*δ*_ is the Huber loss with threshold *δ*:

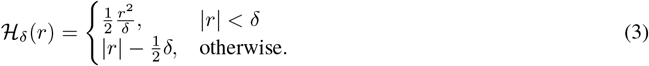

This formulation has two advantages. First, the percentile-based weighting directs more learning capacity toward the low-energy regime that determines early retrieval performance. Second, the Huber form preserves robustness to noisy targets and occasional large residuals, preventing a small number of difficult examples from dominating optimization as they would under a heavily weighted mean squared error objective. Together, these properties make the loss better aligned with the ranking goal, where the priority is to avoid missing the best candidates rather than to finely resolve the relative ordering among many weak binders.

#### Training procedure

Models are trained using the Adam optimizer [25] with a learning rate of 10^*−*4^ and weight decay of 10^*−*5^. Mini-batches of size 20 are used. Gradient norms are clipped to 1.0 to ensure training stability.

A learning rate scheduler (ReduceLROnPlateau) is applied based on validation loss, reducing the learning rate by a factor of 0.5 when performance plateaus.

#### Model selection

Although the model is trained using a regression objective, model selection is performed using ranking-based validation metrics. Specifically, checkpoints are selected based on the mean reciprocal rank (MRR) computed on a held-out validation benchmark. Early stopping is applied with a patience of 10 epochs.

This strategy ensures that the selected model is optimized for the downstream objective of correctly prioritizing true binders rather than minimizing global regression error.

#### Hyperparameter selection

The loss hyperparameters *δ, α*, and *s* were selected by manual grid search over a small set of candidate values evaluated on the held-out validation benchmark using mean reciprocal rank (MRR) as the selection criterion. Specifically, *δ* ∈ {0.01, 0.03, 0.05, 0.10} was evaluated to control the Huber transition point; *α* ∈ {0.1, 0.2, 0.3, 0.5} was evaluated to control the strength of upweighting for strong binders; and *s* ∈ {0.02, 0.05, 0.10 }was evaluated to control the smoothness of the weighting transition. The combination *δ* = 0.03, *α* = 0.3, and *s* = 0.05 yielded the highest validation MRR among all combinations evaluated and was used for all reported experiments. The percentile threshold *p*_40_ was not tuned but computed directly from the empirical 40th percentile of the training-set energy distribution.

Sequence representations are obtained from ESM2 [18], specifically the `esm2_t30_150M_UR50D` checkpoint (30 transformer layers, 150 M parameters, hidden dimension 640). Per-residue embeddings from the final hidden state are extracted by stripping the `[CLS]` and `[EOS]` special tokens, yielding one 640-dimensional vector per residue. These are projected into the shared 128-dimensional structural latent space via a learned linear projection before being fused with atom-level structural features by element-wise summation. During training, all ESM2 parameters are frozen except for the final three transformer encoder layers (layers 27–29), which are fine-tuned jointly with the rest of the model.

### 4.6 Evaluation protocol on unseen antibody–antigen interfaces

#### 4.6.1 Primary ranking benchmark

To evaluate ranking performance in a realistic design scenario, we constructed an independent benchmark consisting of 45 experimentally determined antibody–antigen complexes. For each complex, the CDR3 loop of the antibody and the corresponding interacting epitope residues were extracted from the solved structure.

For every epitope in this benchmark set, we generated 100 candidate paratope sequences using a proprietary inverse folding model (AGZAgen). Sequence generation was conditioned on both the paratope backbone geometry and the target epitope structure and sequence, with the objective of producing sequences capable of folding to the desired backbone conformation while maintaining compatibility with the epitope interface.

For each generated candidate, side-chain conformations were reconstructed using rotamer library–based packing methods to obtain full atomic structures (Fig. 1). The resulting complexes were then scored using the proposed ranking model (AGZArank), which predicts the normalized pseudo-energy learned during training.

Ranking performance was assessed by measuring the ability of a scoring function to retrieve the experimentally observed paratope sequence among the generated candidates. For each target epitope, candidate sequences were ranked according to the model score. Performance was reported as the fraction of cases in which the true sequence appeared within the top-*k* fraction of candidates (1%, 3%, 5%, 10%, 20%, and 50%). In addition to exact sequence recovery, we also evaluated whether the sequence most similar to the experimental paratope was retrieved within the same top-*k* thresholds.

#### 4.6.2 Temporal-shift subset

To assess generalization under distribution shift, we constructed a temporally filtered subset of the primary benchmark consisting of antibody–antigen complexes released after August 2021. Starting from the full set of 45 PDB groups, this filtering resulted in a subset of 15 complexes.

This filtering ensures that evaluated structures were not present in the training data of baseline models such as ProteinMPNN [1] and ESM-IF [2], enabling a more stringent test of generalization to previously unseen interfaces.

All evaluation procedures, including candidate generation, scoring, and ranking metrics, were kept identical to those used in the primary benchmark.

#### 4.6.3 External mutation-based validation (AB-Bind)

To further evaluate biological relevance, we assessed model performance on an external dataset of experimentally measured affinity changes derived from the AB-Bind database [19], accessed via the Natural Antibody resource [26]. This benchmark provides experimentally validated ΔΔ*G* measurements for antibody–antigen complexes with point mutations and is widely used for evaluating mutation-level binding affinity prediction.

We selected this dataset to complement the primary ranking benchmark by testing whether the learned scoring function captures experimentally observed affinity changes, rather than only relative ranking among synthetic or generated candidates.

To ensure compatibility with the model input representation and maintain consistency with prior AB-Bind evaluations, we applied the following filtering criteria: (i) only entries with reported ΔΔ*G* values were retained; (ii) mutations were restricted to the antibody heavy chain; (iii) only mutations within the CDR-H3 region were considered; (iv) complexes were required to have at least five interacting epitope residues, defined as residues containing at least one heavy atom within 4.5 Å of any paratope heavy atom.

After filtering, the dataset was reduced to 96 variants from the original AB-Bind collection, of which 7 correspond to improved-affinity (“tight binder”) mutations. Tight binders were defined using the standard threshold ΔΔ*G <* −0.5 kcal/mol, consistent with the original AB-Bind study.

For each mutation, candidate variants were ranked according to the predicted pseudo-energy, where lower values indicate stronger predicted binding. Performance was evaluated using enrichment-based metrics, measuring the concentration of improved-affinity variants within top-ranked subsets relative to random expectation.

#### 4.6.4 Robustness evaluation

To evaluate robustness to structural perturbations and candidate composition, we performed additional analyses on benchmark variants with controlled geometric perturbations. Specifically, the paratope was subjected to random rigid-body rotation and translation relative to the epitope, together with random coordinate jitter, using the same protocol and parameter ranges as in training-set augmentation. We also evaluated subsets containing only generated (GenAI) candidates, excluding the original experimentally observed sequences. These analyses assess the stability of the ranking function under imperfect structural inputs and altered candidate composition, reflecting practical conditions in generative antibody design workflows.

#### 4.6.5 Baseline comparisons

The proposed ranking model (AGZArank) was compared with two inverse-folding baselines, ProteinMPNN [1] and ESM-IF [2], as well as the structure-based complex affinity predictor PRODIGY [8]. These baselines were selected because prior benchmarking studies have shown that inverse-folding models can provide strong structure-conditioned sequence ranking signals and, in some settings, outperform traditional structure-based scoring approaches such as FoldX for candidate prioritization [11, 12]. This makes ProteinMPNN and ESM-IF particularly relevant comparators for evaluating ranking performance in generative design workflows. PRODIGY was included as a representative affinity-oriented complex scoring method.

Similar to the evaluation protocol used in AbBiBench [11], ProteinMPNN and ESM-IF were not used to generate candidate sequences in this benchmark. Instead, all candidate paratope sequences were taken from the same preconstructed ranking benchmark used for AGZArank evaluation, and the inverse-folding baselines, as well as PRODIGY, were used only to rescore these candidates for reranking.

For the inverse-folding baselines, the epitope chain was treated as the fixed structural context, whereas the paratope chain was treated as the designed chain. Each benchmark group consisted of one experimentally observed paratope–epitope pair together with 100 generated candidate paratope sequences. Given the corresponding paratope–epitope fragment structure, ProteinMPNN and ESM-IF were used to compute the conditional log-likelihood of each candidate paratope sequence under this fixed/designed chain assignment, thereby measuring how compatible the candidate sequence was with the paratope backbone in the presence of the fixed epitope context. For ProteinMPNN, we used the official pretrained `v_48_020` model with vanilla weights. For ESM-IF, we used the pretrained `esm_if1_gvp4_t16_142M_UR50` checkpoint. The resulting log-likelihood was used as the ranking signal, with higher values indicating better-ranked candidates.

For PRODIGY, affinity was estimated from antigen–paratope complexes composed of the original full antigen chain and the candidate paratope structure, with the paratope rebuilt from the candidate sequence and backbone as described for the evaluation benchmark. Candidates were ranked by predicted affinity, with more favorable predicted binding ranked higher.

## Acknowledgments

The authors acknowledge support from the NVIDIA Inception Program and the StartX accelerator program, which provided access to AWS cloud computing credits through AWS Activate. These credits were used to support model development and evaluation in this study.

## Author Contributions

D.S. defined the task formulation, developed the overall modeling strategy, designed the training data, benchmark, and evaluation strategy, shaped the scientific framing of the manuscript and supervised the project. Z.S. contributed to model design, implemented the model and training pipeline, performed the experiments, analyzed the results and wrote the initial manuscript draft. A.K. contributed to model development by implementing and evaluating alternative architectural directions and participated in experimentation. B.S. contributed biological insight, participated in data and evaluation design, tested alternative architectural directions and provided strategic guidance. All authors discussed the results, reviewed and revised the manuscript and approved the final version.

## Competing interests

This work is related to a pending non-provisional patent application assigned to Arlan Biotech Inc. All authors are named as inventors on the application. D.S. and B.S. are shareholders of Arlan Biotech Inc.; Z.S. and A.K. are employees of Arlan Biotech Inc.

## Code availability

The AGZArank model implementation is available at: https://github.com/jenisbek-git/AGZArank. The repository includes training, inference, and evaluation pipelines, along with instructions for reproducing the results reported in this study.

## Data availability

The benchmark data developed in this study are available at: https://zenodo.org/records/20581998. The Zenodo record includes the benchmark test set in PDB format, source data, baseline inputs and outputs, instructions for reproducing the ESM-IF, ProteinMPNN, and PRODIGY results, Graphinity enrichment derivation files, and AB-Bind preparation files for AGZArank inference.

The AGZArank benchmark output files used to generate the reported tables are available in the associated GitHub repository. The synthetic dataset used to train AGZArank and the associated production model checkpoints are not publicly released because they are proprietary components of Arlan Biotech Inc.’s binder-design platform. Access to AGZArank checkpoint-based inference may be provided by Arlan Biotech Inc. upon reasonable request.

## Notes

https://github.com/jenisbek-git/AGZArank

https://zenodo.org/records/20581998

